# P-KNN: Maximizing variant classification evidence through joint calibration of multiple pathogenicity prediction tools

**DOI:** 10.1101/2025.09.24.678417

**Authors:** Po-Yu Lin, Nadav Brandes

## Abstract

Clinical guidelines for Mendelian disease diagnosis require that outputs from variant pathogenicity prediction tools be converted into well-calibrated probabilities. However, the existing calibration process is only valid when pre-committing to a specific tool, preventing clinicians from using multiple tools with complementary strengths. To lift this restriction, we introduce Pathogenicity K-Nearest Neighbors (P-KNN), a simple, flexible method that jointly calibrates any set of tools. P-KNN scores each variant by the fraction of pathogenic variants among those with the most similar scores across all underlying tools. We compared P-KNN to single-tool calibration over 13 real tools, including two meta-predictors. Compared to BayesDel, the best-performing tool, P-KNN produced better-calibrated probabilities and stronger evidence strength, with mean log likelihood ratios of 2.56 vs. 2.24 for pathogenic variants, 2.29 vs. 1.56 for benign variants, and 2.28 vs. 2.07 for variants of uncertain significance. We also evaluated P-KNN at four historical time points to assess scalability and robustness, finding that it improved steadily as new tools became available, while remaining well-calibrated. It also correctly integrated computational predictions with experimental measurements, whereas guidelines-based evidence summation systematically overestimated pathogenicity. In summary, P-KNN allows full flexibility to work with any set of tools in a reliable and robust manner. It is consistent with clinical variant classification guidelines, making it well positioned to improve diagnostic yield for rare genetic diseases. P-KNN is available via command line (https://github.com/Brandes-Lab/P-KNN) and precomputed scores (Dataset: https://huggingface.co/datasets/brandeslab/P-KNN, User Interface: https://huggingface.co/spaces/brandeslab/P-KNN-Viewer).

## INTRODUCTION

Variant classification is the process of determining whether genetic variants cause human monogenic disease. In clinical practice, variant classification is guided by the joint guidelines of the American College of Medical Genetics and Genomics (ACMG) and the Association for Molecular Pathology (AMP) ^1^. The ACMG/AMP guidelines classify variants into one of five categories: pathogenic, likely pathogenic, variant of uncertain significance (VUS), likely benign, or benign. Each category corresponds to a probability that the variant is causally related to a disease (**Figure 1A**). These probabilities are estimated from diverse sources of evidence, including segregation analysis from pedigrees, functional experiments, and computational pathogenicity prediction tools. Subsequent refinements by the ClinGen Sequence Variant Interpretation Working Group have restructured the ACMG/AMP guidelines into a Bayesian framework ^2,3^. In this framework, each source of evidence is quantified as a log-likelihood ratio (LLR) and categorized into 4 levels: supporting (+1), moderate (+2), strong (+4), and very strong (+8). These LLRs are then summed to compute the posterior pathogenicity probability of a given variant, which determines its classification.

**Figure 1:**
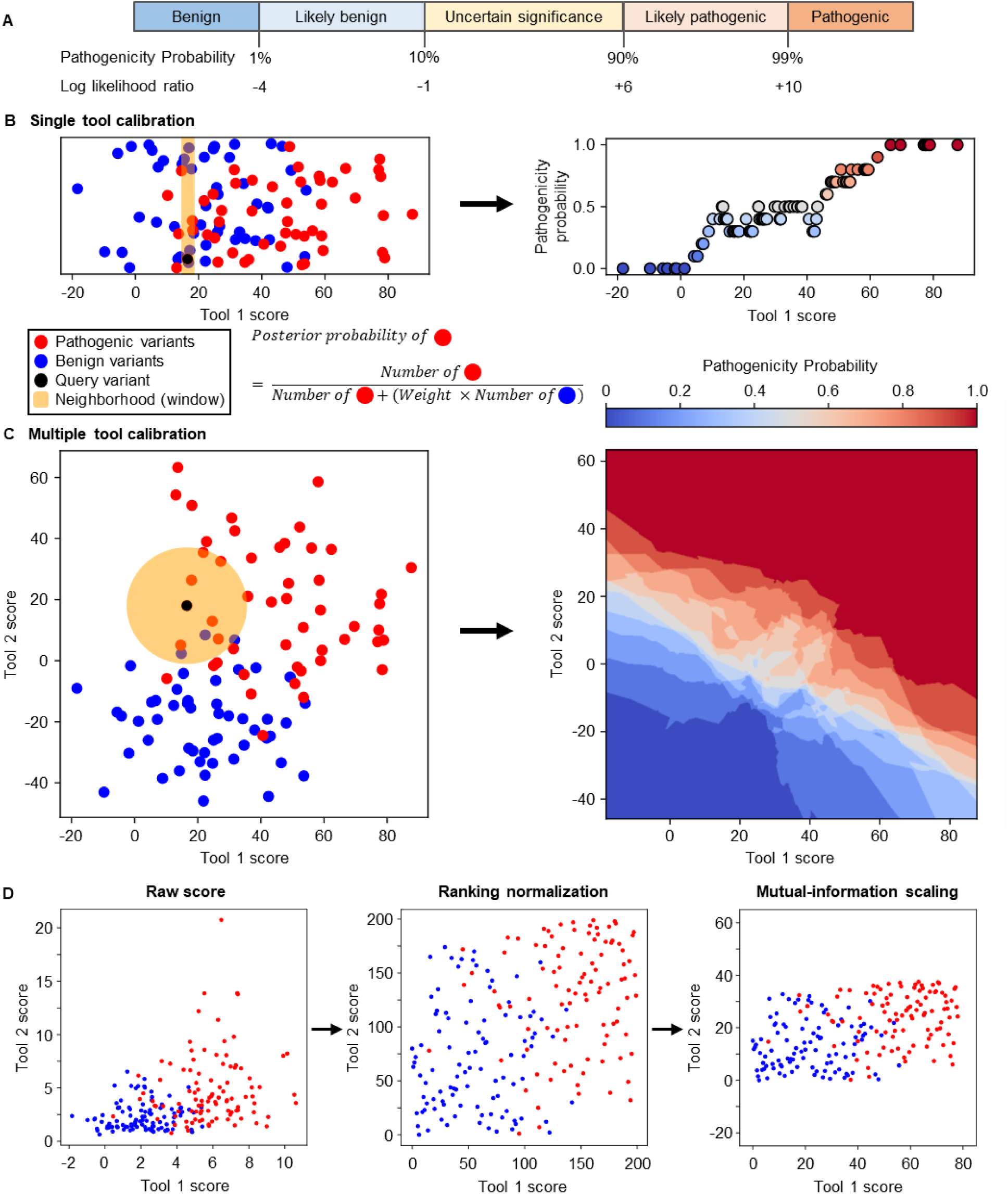
Joint calibration of multiple pathogenicity prediction tools with P-KNN. **(A)** Variant classification categories according to the ACMG/AMP guidelines, with corresponding pathogenicity probabilities and required evidence strength quantified in log likelihood ratio. **(B)** Single-tool calibration operates in one dimension, applying a window to include the variants with the closest scores to the query variant. The proportion of pathogenic variants within the window estimates pathogenicity probability. **(C)** In contrast, P-KNN operates in a high-dimensional space, where each dimension represents the scores from a different tool. A spherical neighborhood is defined around the query variant’s coordinates, and the proportion of pathogenic variants within this neighborhood estimates the probability that the query variant is pathogenic. **(D)** Because tools produce scores on different scales, normalization (e.g., ranking) is performed prior to distance calculation. Each tool is further scaled according to its mutual information with pathogenicity labels, giving more weight to informative tools and reducing the influence of less predictive ones.

In the original ACMG/AMP guidelines, computational pathogenicity prediction tools are only permitted to contribute supporting-level evidence ^1^. However, recent work by ClinGen has demonstrated that, within the Bayesian framework, certain tools can achieve strong levels of evidence ^4,5^. The incorporation of computational tools within this framework requires proper calibration to convert their raw prediction scores into LLRs. The current calibration method proposed by ClinGen is limited to calibrating one tool at a time ^4,5^. This presents a limitation, as different tools often perform better for different genes or clinical phenotypes ^6,7^. Despite this variability, existing guidelines offer no recommendations for tool selection, and the one-tool-at-a-time framework prevents leveraging complementary strengths across tools. Moreover, the guidelines actively caution against trying multiple tools for classifying a given variant, and recommend choosing a tool before looking at any predictions ^5^. Although it is important statistical advice, adhering to this guideline is exceedingly difficult and puts clinicians at a disadvantage of being unable to revise their selection.

An alternative strategy involves training a meta-predictor that integrates multiple tools, followed by calibration of its output scores. While appealing in principle, this approach faces key practical challenges. Calibration requires a reference dataset of labeled pathogenic and benign variants. Yet many variants have already been used in training either the meta-predictor itself or its underlying component tools, which prevents their reuse in calibration ^8^. This substantially reduces the data available for reliable calibration. Moreover, once trained, meta-predictors are rarely updated to incorporate newly developed tools, as doing so typically requires retraining the model from scratch.

To address these challenges, we introduce Pathogenicity K-Nearest Neighbors (P-KNN), an inherently calibrated meta-prediction framework that enables the integration and joint calibration of any set of pathogenicity prediction tools. P-KNN leverages the complementary strengths of individual tools to provide stronger and more reliable evidence than standard single-tool calibration, while remaining robust to the incorporation of newly developed tools. It facilitates dynamic, up-to-date variant classification in the rapidly evolving landscape of computational pathogenicity prediction. With GPU acceleration, P-KNN can calibrate hundreds of query variants per minute. It is available via command-line interface, and precomputed scores are provided for all possible missense variants in the human genome (see Implementation and code availability, Availability of data and materials).

## METHODS

### Overview of the P-KNN joint calibration framework

P-KNN estimates the pathogenicity probability of a genetic variant given output scores from multiple prediction tools. P-KNN requires three input datasets: i) query variants to be calibrated, ii) a calibration dataset containing labeled pathogenic and benign variants, and iii) an unlabeled regularization dataset that reflects the general distribution of variants in the human genome. Following a similar dataset preparation strategy to that of ClinGen, we split ClinVar into a calibration dataset and a held-out test set of query variants ^4,5,9^. We used gnomAD for the regularization dataset ^10^. All three variant datasets must be scored by the same set of tools, though our framework can tolerate missing scores for some of the variants.

P-KNN can be seen as an extension of the standard single-tool calibration framework developed by ClinGen ^4,5^. In single-tool calibration, the scores from a single tool are used to identify the variants in the calibration dataset whose scores are closest to the query variant (**Figure 1B**) ^5^. The pathogenicity probability is then estimated as the proportion of pathogenic variants among these nearest variants. P-KNN extends this principle by representing each variant as a point in a multidimensional space, where each tool’s scores define a dimension. Using the same nearest-neighbor approach, P-KNN identifies the variants within the calibration dataset that are closest to the query variant in this multidimensional space. It then estimates the probability of the query variant being pathogenic as the proportion of pathogenic variants within these nearest neighbors (**Figure 1C**).

### Score standardization

Because the scores produced by different pathogenicity prediction tools may be on entirely different scales, they first need to be normalized (**Figure 1D**). In addition, many tools do not provide genome-wide predictions, resulting in missing values for some variants in the dataset. A strategy for handling missing values is therefore required. Furthermore, since individual tools differ in their ability to discriminate between pathogenic and benign variants, it is important to assign more weight to more informative tools when calculating distances and searching for the nearest neighbors.

#### Score normalization

We evaluated two score normalization methods: ranking and z-score. In ranking normalization, the scores for variants in the calibration dataset produced by each tool are ranked in ascending order. Variants with identical scores are assigned the average of their ranks. Missing values are retained without modification. Variants in the query and regularization datasets are assigned a rank score by interpolating between the variants with immediately lower and higher raw scores in the calibration dataset. In z-score normalization, the arithmetic mean and standard deviation of non-missing scores within the calibration dataset are calculated for each tool. These statistics are then used to perform z-score transformation on the scores from the calibration, regularization and query datasets.

#### Handling missing values

We evaluated two approaches to handle missing scores within the calibration dataset: exclusion (removing variants with any missing score) and score imputation. For the query and regularization datasets, we consistently applied score imputation. We imputed missing scores using KNN imputation ^11^ (not to be confused with P-KNN itself). This algorithm identifies the *k* scored variants in the calibration dataset that are nearest to the missing-score variant, based on Euclidean distance over all available tool scores. To ensure comparability of distance across variants with different numbers of available scores, the computed distance is divided by the square root of the number of tools used. The missing score is then imputed as the average of the missing tool’s scores among these nearest neighbors. The hyperparameter *k* is set to match the minimum number of nearest neighbors used for pathogenicity probability estimation (*N*_*calibration*_= 100 defined in “Posterior pathogenicity probability calculation”).

#### Mutual-information scaling

We scaled the scores from each tool by their mutual information with pathogenicity labels within the calibration dataset, thereby prioritizing more informative tools and reducing the influence of low-performing ones. To estimate the mutual information between the discrete pathogenicity label and the continuous score of a given tool, we employed a non-parametric KNN estimator specifically designed for discrete-continuous variable pairs ^12,13^ (Supplementary Methods, mutual-information scaling).

### Posterior pathogenicity probability calculation

Following score standardization, the next steps are to i) identify the nearest variants for each query variant within the calibration dataset, and ii) estimate the pathogenicity probability as the weighted proportion of pathogenic variants among these nearest variants ^14^. These two steps follow the same logic as in the standard single-tool calibration framework defined by ClinGen ^5^.

#### Identify the nearest neighbors of each query variant

To define the number of nearest neighbors for each query variant, we introduced two hyperparameters: *N*_*calibration*_and *N*_*regularization*_. These hyperparameters specify the minimum number of variants from the calibration and regularization datasets, respectively, that must be included within the spherical neighborhood used for probability estimation. For each query variant, we compute the Euclidean distance to all variants in both the calibration and regularization datasets. We then identify the distances to the *N*_*calibration*_th nearest neighbor in the calibration dataset and to the *N*_*regularization*_th nearest neighbor in the regularization dataset. The final search radius is defined as the larger of these two distances. All variants within the calibration dataset falling within this radius are selected as the nearest neighbors used for pathogenicity probability estimation. Following the same settings defined by ClinGen for single-tool calibration, *N*_*calibration*_ is set to 100, and *N*_*regularization*_ is set to 3% of the total number of variants in the regularization dataset (genomAD) ^5^.

#### Calculate the proportion of pathogenic variants

Let *N*_*pathogenic*_ and *N*_*benign*_ denote the total numbers of pathogenic and benign variants in the calibration dataset, respectively, and let *n*_*pathogenic*_and *n*_*benign*_represent the corresponding counts among the chosen nearest neighbors of a given query variant. We first compute a weighting factor *w* that aligns the global pathogenic-to-benign ratio in the calibration dataset with a predefined prior pathogenicity probability *P*_*prior*_. This is achieved by solving *P*_*prior*_ = 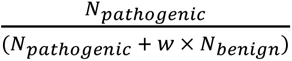. This weight *w* is then applied to the local neighbor counts to estimate the posterior pathogenicity probability for the query variant as a weighted proportion *P*_*post*_ = 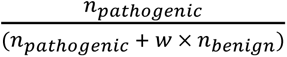. We set *P*_*prior*_ to 0.0441, following the value recommended by ClinGen, which was derived from a large-scale investigation of gnomAD using the DistCurve algorithm ^5,10,15^.

### Bootstrapping and calculating posterior probabilities

To improve the robustness of posterior probability estimation, we apply the same bootstrapping process defined by ClinGen ^5^. As part of this bootstrapping process, the entire pipeline, including score standardization and posterior pathogenicity probability calculation, is repeated across multiple resamples of the calibration dataset. Each resample contains the same number of variants as the original calibration set, sampled with replacement. Each iteration yields a posterior pathogenicity probability *P*_*post*_ for the query variant, with the corresponding benignity probability 1 − *P*_*post*_. After completing all bootstrap iterations, we compute one-sided 95% confidence intervals. The P-KNN posterior probability for the query variant being pathogenic is defined as the 5th percentile of the obtained *P*_*post*_distribution, and the posterior probability for it being benign is defined as the 5th percentile of the 1 − *P*_*post*_distribution. These provide conservative estimates for the posterior probability of a variant being pathogenic or benign. Because both values are taken as the 5th percentiles of their respective distributions, their sum is typically less than 100%, reflecting the intentional conservatism of this estimation approach.

### Conversion of posterior probabilities to log likelihood ratios measuring evidence strength

We followed ClinGen’s procedures to convert posterior probabilities into LLRs that quantify the strength of evidence supporting a variant as being pathogenic or benign within ACMG/AMP Bayesian framework ^2,3^. LLRs are calculated using the standard formula for a Bayesian update of the log odds, 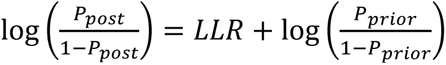.We apply this formula separately for each of the two pathogenicity labels to calculate *LLR*_*pathogenic*_ and *LLR*_*benign*_, each using the corresponding prior and posterior probabilities calculated previously (following bootstrapping). The choice of the logarithmic base is based on the same method specified by ClinGen ^2,3^. The resulting LLRs are clipped to a range of [0, 8] to ensure compatibility with the ACMG/AMP Bayesian framework. This ensures that at least one of *LLR*_*pathogenic*_ and *LLR*_*benign*_is always 0. Finally, the overall evidence strength in support of a variant being pathogenic is calculated as *LLR*_*overall*_ = *LLR*_*pathogenic*_ − *LLR*_*benign*_. This measure of evidence strength is fully compatible with the ACMG/AMP Bayesian framework and can be directly applied in clinical variant classification.

### Evaluating P-KNN and single-tool calibration

We evaluated P-KNN and single-tool calibration over a held-out test set from ClinVar consisting of query variants with known pathogenicity labels ^9^. Performance was assessed based on two metrics: mean evidence strength and calibration.

#### Mean evidence strength

We computed the mean *LLR*_*overall*_ separately for pathogenic and benign variants in the test set. To ensure a uniform direction, the mean for benign variants was multiplied by -1. In some analyses, we averaged the mean *LLR*_*overall*_across pathogenic variants and across benign variants (assigning a weight of ½ to each of the two groups), in order to produce a single score reflecting the ability of P-KNN or individual tools to assign strong and directionally correct evidence across the two variant classes.

#### Calibration

Calibration assesses how well the calibrated posterior probabilities from P-KNN or single-tool calibration align with the true frequencies of pathogenic and benign variants in the test set. For posterior probabilities of being pathogenic, the [0, 1] range of posterior probabilities is divided into 10 equal bins. Within each bin, we count the number of pathogenic and benign variants, *n*_*pathogenic*_ and *n*_*benign*_, in the test set that are assigned posterior probabilities within this range. We then estimate the bin-specific frequency of pathogenic variants *f*_*pathogenic*_ as the weighted proportion of pathogenic variants 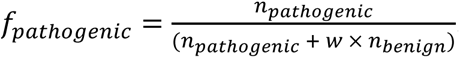. The weighting factor *w* aligns the global pathogenic-to-benign ratio in the test set with the predefined *P*_*prior*_ (see “Posterior pathogenicity probability calculation”). To quantify uncertainty, we construct a two-sided 95% confidence interval for the number of pathogenic variants in each bin using a binomial distribution. These boundary counts are then transformed into confidence intervals for *f*_*pathogenic*_. If the upper bound of the confidence interval for the true frequency of pathogenic variants within a bin falls below the posterior probability defined by that bin, we consider the posterior probabilities in that range to be miscalibrated. The same procedure is repeated for posterior probabilities of being benign, except that the evaluation is limited to the [0.95, 1.0] range of posterior probabilities, because only in this range the posterior probability exceeds the prior probability of being benign.

### Implementation and code availability

We developed two independent code bases: one optimized for accelerated GPU computing, and the other for multi-CPU parallelization. The GPU implementation utilizes PyTorch, while the CPU implementation is built on NumPy ^16,17^. P-KNN can be run via a command-line interface available at https://github.com/Brandes-Lab/P-KNN. In addition, precomputed scores based on 27 tools in dbNSFPv5.2a are available at https://huggingface.co/datasets/brandeslab/P-KNN ^18^.

## RESULTS

### Simulation analyses identify ranking normalization and mutual-information scaling as the most robust score standardization

Since P-KNN is sensitive to the scaling of scores from the underlying variant pathogenicity prediction tools, we sought to identify an optimal score standardization pipeline that would provide strong and robust predictions. We created a simulated dataset of pathogenic and benign variants and trained 50 different pathogenicity prediction tools on this simulated data (Supplementary methods, Evaluation with simulated tools). We evaluated four score standardization pipelines by varying two factors: normalization method (ranking vs. z-score) and the inclusion or exclusion of mutual-information scaling. We then applied P-KNN on subsets of the 50 underlying tools, using each of the four parameter combinations (**Figure 2A-B**). We observed that the combination of ranking normalization and mutual-information scaling yielded the strongest evidence across varying numbers of underlying tools and fractions of underlying tools that are highly informative. We fixed the normalization and scaling strategy to this optimal combination in all subsequent analyses.

**Figure 2.**
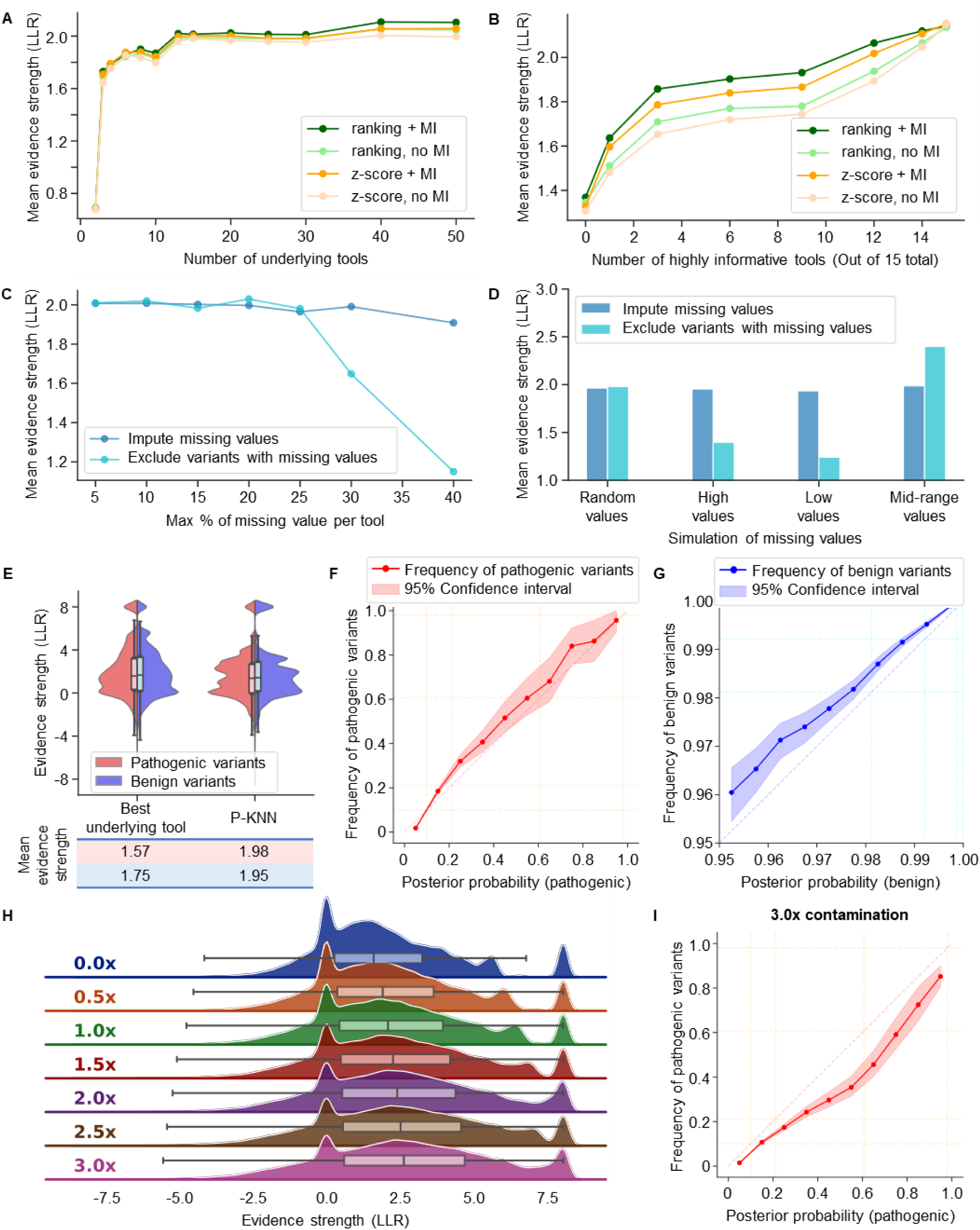
Evaluation of P-KNN on simulated pathogenicity prediction tools. **(A, B)** Mean evidence strength (log-likelihood ratio, LLR) estimated by P-KNN under different normalization methods (ranking vs. z-score) and inclusion or exclusion of mutual-information scaling. **(A)** As a function of the number of integrated tools. **(B)** As a function of the number of highly informative tools (out of 15 total). **(C, D)** Mean evidence strength (LLR) under different strategies for handling missing tool scores: imputation vs. exclusion. **(C)** As a function of the maximum percentage of missing scores per tool (each tool assigned a random missing rate between 0 and the maximum; missing values inserted randomly). **(D)** Under different missingness patterns (random, or biased toward high, low, or mid-range scores, with a maximum missing rate of 25%). **(E)** Evidence strength of P-KNN (over 15 randomly selected tools) compared to the best-performing underlying tool. **(F, G)** Calibrated posterior probabilities from P-KNN vs. actual frequencies of pathogenic/benign variants. **(H)** Change in evidence strength with increasing amount of data contamination between the training and calibration datasets (e.g., 3.0x indicates that the calibration dataset includes 3 variants from the training set of the underlying tools for every variant in the original, held-out calibration dataset). **(I)** High degree of data contamination (3.0x) leads to inflated posterior probabilities.

### KNN imputation best handles missing values in simulations

Since many computational prediction tools do not provide genome-wide predictions, missing scores are inevitable. We evaluated two approaches for handling missing values in the calibration dataset: imputation or exclusion of variants with missing values. We randomly selected 15 simulated tools and deleted some of the scores within each of these tools. As the percentage of missing values increased, imputation preserved evidence strength better than variant exclusion (**Figure 2C**). Furthermore, when missing values are not uniformly distributed across all variants, but are instead biased to higher or lower score ranges, imputation maintains greater stability (**Figure 2D**). Based on these results, we selected to impute missing values in all subsequent analyses.

### P-KNN is well-calibrated and outperforms the underlying simulated tools

We evaluated the complete score standardization pipeline, comprising ranking normalization, imputation, and mutual-information scaling, using 15 randomly selected simulated tools, each with up to 25% missing values. P-KNN produced stronger evidence for both pathogenic and benign variants than all the underlying tools calibrated individually (**Figure 2E**). In addition, the posterior probabilities are almost perfectly calibrated (**Figure 2F-G**).

### Contamination of the calibration dataset leads to miscalibration

We also investigated how data contamination affects P-KNN by progressively increasing the overlap between the calibration dataset and the training data of the underlying tools. We used the same randomly selected 15 simulated tools, each containing up to 25% missing values. In addition to the original variants in the held-out calibration dataset, we incrementally added variants from the training set of the underlying tools into the calibration dataset. As the degree of contamination increased, we observed an overestimation of evidence strength (**Figure 2H**), along with corresponding miscalibration (**Figure 2I**).

### P-KNN gains more evidence from established tools on real clinical data

Following the promising simulation results, we sought to test P-KNN over real variant pathogenicity prediction tools and data. To this end, we considered the same 13 tools and dataset used in the original ClinGen study establishing the single-tool calibration framework ^5^. This dataset took pathogenic and benign variants reported to ClinVar before December 2019 as the calibration dataset, pathogenic and benign variants reported to ClinVar in 2020 as the test set, and gnomAD as the regularization dataset ^9,10^ (Supplementary methods, Evaluation with established tools on real clinical data). We applied P-KNN to jointly calibrate all 13 tools, and compared its performance against singe-tool calibration of each of the underlying tools. Compared to the four best-performing tools (BayesDel ^19^, REVEL ^20^, MutPred2 ^21^ and VEST4 ^22^), P-KNN assigned stronger directionally-correct evidence for both pathogenic and benign variants (**Figure 3A**). Specifically, compared to BayesDel, the top-performing tool, P-KNN produced higher mean evidence strength for both pathogenic (2.56 vs. 2.24, p=5E-54) and benign variants (2.29 vs. 1.56, p<1E-100). P-KNN also assigned directionally-correct evidence in support of pathogenicity to a higher percentage of pathogenic variants (78.8%), compared to BayesDel (74.7%), REVEL (71.9%), MutPred2 (73.6%), and VEST4 (74.1%). Similarly, it assigned directionally-correct evidence to the highest percentage of benign variants (75.1%), compared to BayesDel (68.2%), REVEL (66.4%), MutPred2 (70.8%) and VEST4 (67.7%) (**Supplementary Figure 1**).

**Figure 3.**
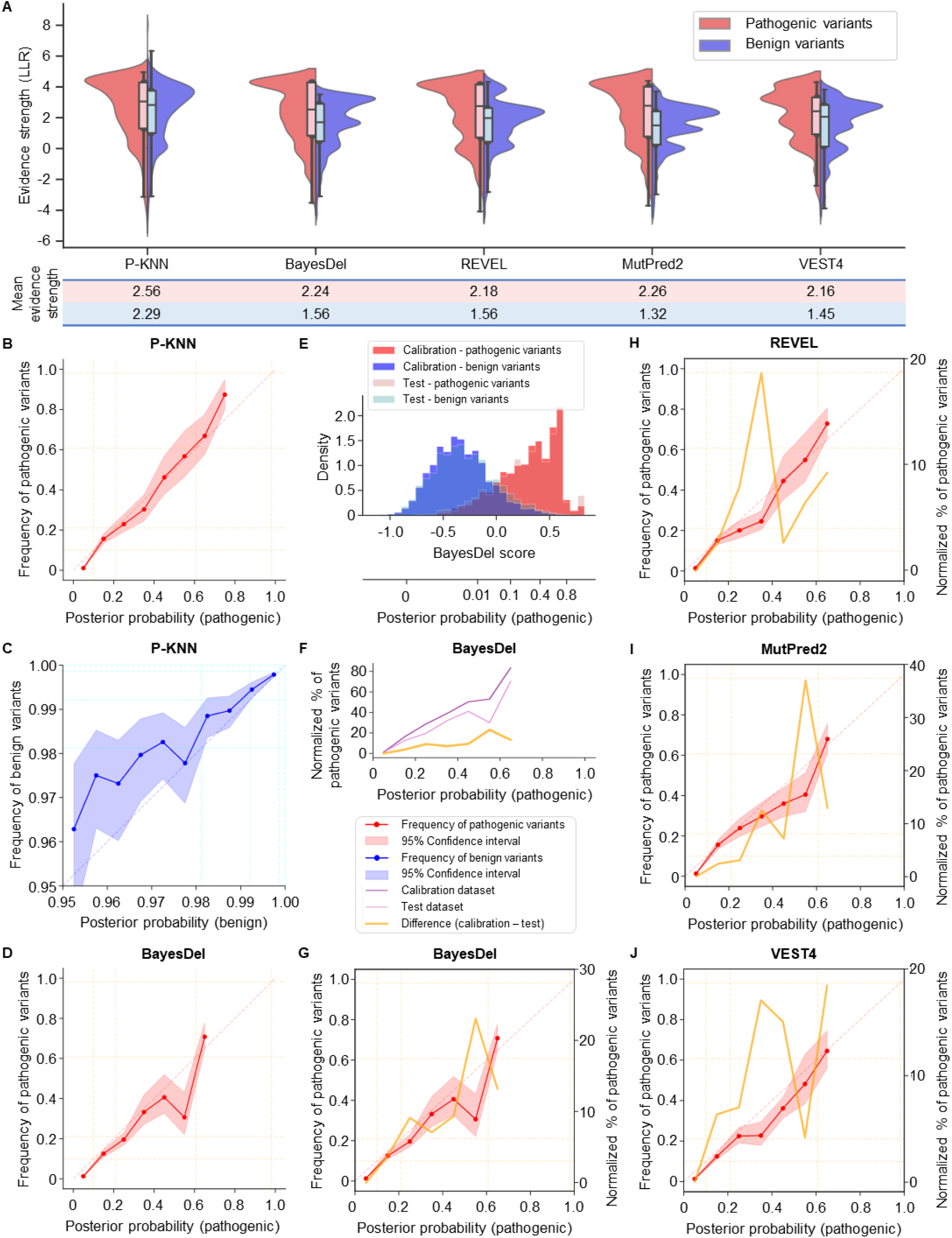
Evaluation of P-KNN over individual tools. **(A)** Evidence strength (log likelihood ratio, LLR) of P-KNN integrating 13 commonly-used variant effect prediction tools, and the four best-performing individual tools, over pathogenic and benign variants. **(B-C)** Calibration of P-KNN on the held-out test set (as in Fig. 2F-G). **(D)** Single-tool calibration of BayesDel, showing notable miscalibration of posterior probabilities in the 0.5-0.6 range. **(E)** The distributions of the raw BayesDel prediction scores (and the posterior probabilities they are converted to post calibration) across pathogenic and benign variants in the calibration and test datasets. **(F)** The normalized percentage of pathogenic variant as a function of posterior probabilities from BayesDel in the calibration dataset, test dataset, and the difference between them. **(G-J)** Overlaying the calibration curve (D) alongside the differences in normalized percentages of pathogenic variants between the calibration and test datasets (F) for BayesDel, REVEL, MutPred2, and VEST4. The score ranges with different percentages of pathogenic variants coincide with the regions of miscalibration.

### P-KNN is well-calibrated, while single-tool calibration shows miscalibration

P-KNN also provides reliable, well-calibrated posterior probabilities, consistent with the true frequencies of pathogenic and benign variants in the held-out test set (**Figure 3B-C**). In contrast, miscalibration was frequent in single-tool calibrations. For example, BayesDel shows miscalibration in the 0.5–0.6 posterior probability range (**Figure 3D**). To the best of our knowledge, the miscalibration of the standard single-tool calibration framework over prominent pathogenicity prediction tools has not been previously reported. Given the significance of this finding to variant classification and genetic diagnosis, we sought to understand its root cause. We found that miscalibration arises from discrepancies in the score distributions of pathogenic and benign variants between the calibration and test datasets (**Figure 3E**), which consist of variants submitted to ClinVar before 2019 and during 2020, respectively ^5,9^. To highlight this distribution shift, we binned variants by their posterior probabilities and examined the proportion of pathogenic variants in the calibration and test dataset within each bin (**Figure 3F**). We then overlayed the differences in pathogenicity frequencies between the two datasets alongside the calibration curve (**Figure 3G**). This analysis reveals that score ranges showing miscalibration correspond to regions with distribution shift between the two datasets. Similar patterns were observed for REVEL, MutPred2, and VEST4 (**Figure 3H–J**).

### P-KNN provides stronger evidence for variants of uncertain significance

Next, we sought to investigate how the improved evidence from P-KNN translates to improved classification of VUS, which are a critical major challenge in clinical genetics. To this end, we applied P-KNN to jointly calibrate 11 tools over VUS in ClinVar. Although the full ClinGen dataset includes 13 prediction tools, only 11 of them were available in the dbNSFP database, which served as the source for prediction scores for VUS ^18^. Compared to BayesDel, the top-performing among these 11 tools, P-KNN achieved a significantly higher mean evidence strength for VUS (2.28 vs. 1.94, p<1E-100). P-KNN exceeded supporting-level evidence (|LLR| > 1) over a greater proportion of VUS (75.2%) than the three best-performing single tools: BayesDel (74.5%), REVEL (68.8%), and VEST4 (70.1%) (**Supplementary Figure 2**). At the same time, BayesDel and VEST4 assigned more VUS with evidence in support of being pathogenic than P-KNN did, but this is likely due to their tendency to assign a larger fraction of benign variants with incorrect evidence for being pathogenic (Supplementary Figure 1).

### P-KNN outperforms and benefits from supervised meta-predictors

P-KNN can be viewed as an inherently calibrated meta-predictor: it takes predictions from multiple tools as inputs and outputs a single score that consolidates the underlying predictions. A natural question is how P-KNN compares to the traditional, supervised meta-predictors. To answer this question, we compared P-KNN against two state-of-the-art meta-predictors: REVEL ^20^ and BayesDel ^19^ (Supplementary method, P-KNN vs. meta-predictors). To ensure fair comparison, we applied P-KNN over the same 18 component tools that REVEL uses as inputs (BayesDel was trained on a subset of these tools). Compared to single-tool calibration of REVEL, P-KNN produces overall stronger evidence, with slightly lower evidence strength for pathogenic variants (mean LLR of 2.10 vs. 2.18) but substantially higher for benign variants (1.99 vs. 1.56; **Figure 4A**). When REVEL’s prediction scores were included as an additional input to P-KNN, evidence strength increased for both classes (mean LLRs of 2.18 and 2.03 for pathogenic and benign variants, respectively). Incorporating BayesDel led to further gains in evidence strength (mean LLRs of 2.25 and 2.15 for pathogenic and benign variants, respectively), outperforming both REVEL (2.18 and 1.56) and BayesDel (2.24 and 1.56). While single-tool calibration of REVEL and BayesDel exhibited miscalibration, all three variations of P-KNN demonstrated consistently good calibration (**Figure 4B**). These results suggest that P-KNN not only effectively integrates multiple tools but also benefits from incorporating meta-predictors using the same component tools, enhancing evidence strength while preserving robust calibration.

**Figure 4.**
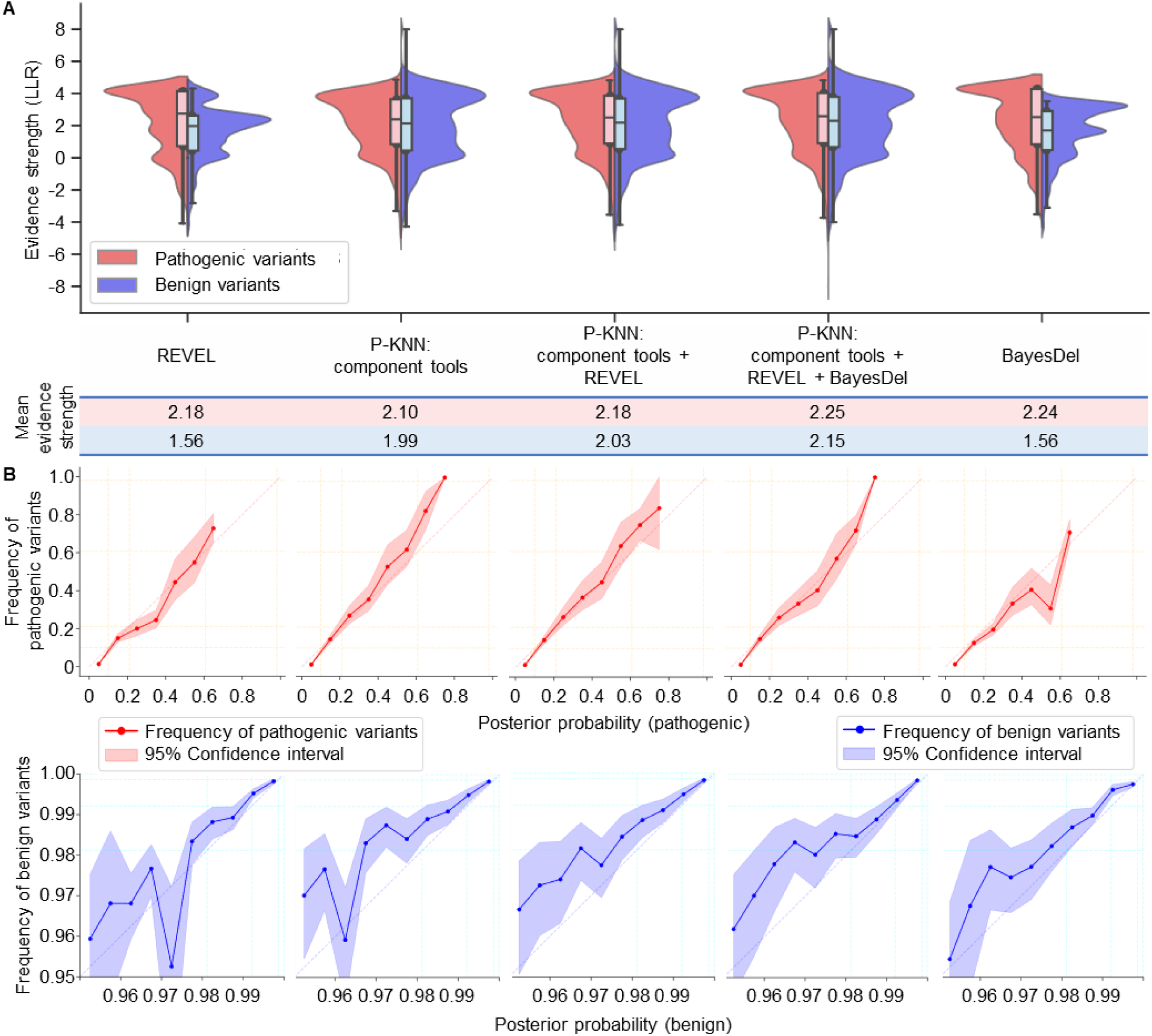
Comparison between P-KNN and supervised meta-predictors. **(A)** Evidence strength (log likelihood ratio, LLR) obtained by i) single-tool calibration of REVEL, ii) P-KNN applied to the 18 component tools of REVEL, iii) P-KNN over the same component tools and REVEL itself, iv) P-KNN additionally applied over BayesDel, and v) single-tool calibration of BayesDel. (B) Calibration curves for the above methods.

### P-KNN consistently improves through integration of newer prediction tools

With the ongoing introduction of new computational pathogenicity prediction tools with improved predictions, a natural question is whether P-KNN can repeatedly incorporate new tools in order to remain up to date. To evaluate how P-KNN performance would have evolved over time, we sequentially incorporated tools released at four historical time points based on different versions of the dbNSFP database ^18,23^. After excluding tools with known or potential overlap training data with the calibration dataset, we applied P-KNN to the remaining tool sets, which comprised 18, 22, 27 and 27 tools, respectively (Supplementary method, P-KNN evolution with new tools). The details of all included tools are available in **Supplementary Table 1**.

As newer tools were added, the mean evidence strength produced by P-KNN progressively increased (**Figure 5A**) while remaining consistently well-calibrated (**Figure 5B**). Compared to AlphaMissense, the best-performing single tool, P-KNN incorporating the newest set of tools assigned overall stronger evidence, with slightly lower mean LLR for pathogenic variants (2.73 vs. 2.85) but substantially higher for benign variants (2.31 vs. 1.58) ^24^. These findings indicate that, provided that the calibration dataset is free from contamination by variants used in the training of the newly introduced tools, P-KNN can robustly incorporate and benefit from advancements in pathogenicity prediction tools.

**Figure 5:**
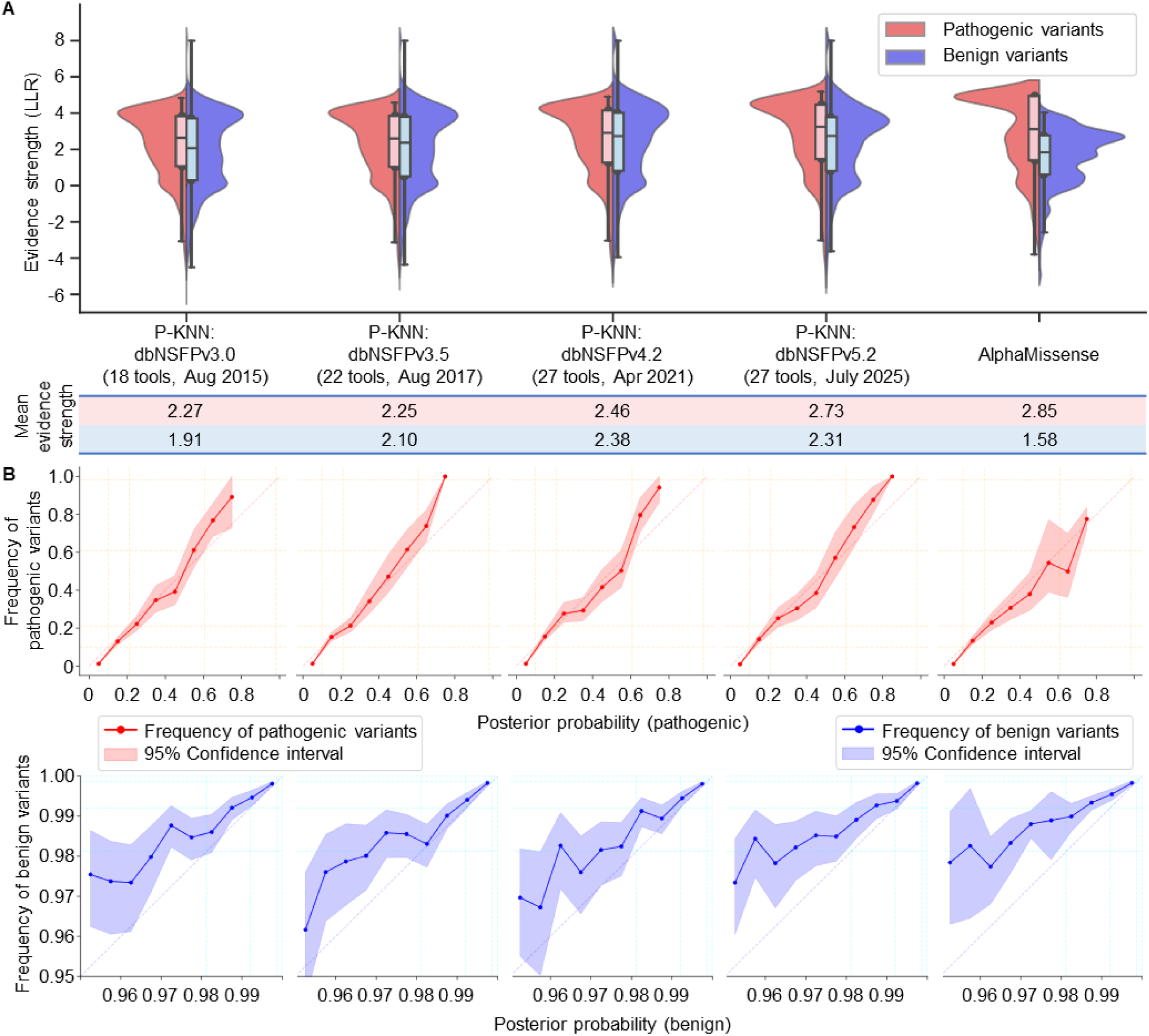
Applying P-KNN over tool sets from four historical time points. **(A)** Evidence strength (log likelihood ratio, LLR) obtained by applying P-KNN over the sets of tools available at four historical time points, alongside AlphaMissense (the best-performing among all the individual tools). **(B)** Calibration curves for the above methods.

### Experimental and computational evidence are correlated and require joint calibration

Deep mutational scanning (DMS) assays have emerged as a powerful source of variant-level measurements of functional genetic effects, with increasing efforts dedicated to calibrating these scores ^25–27^. Within the ACMG/AMP Bayesian framework, DMS are categorized as functional evidence, while computational scores are predicted evidence ^1,2^. The LLRs derived from these two sources are typically combined by simple addition under the assumption of statistical independence. However, if this assumption does not hold, joint calibration of these distinct evidence types may be necessary.

To evaluate the necessity of joint calibration, we examined the 324 *TP53* variants with clear pathogenic/benign labels in ClinVar ^9^. These variants were annotated with 16 DMS assays, including one assessing dominant-negative effects (DMS_domneg) ^28^ and another measuring transcriptional activity (DMS_txact) ^29^. As a representative computational pathogenicity prediction tool, we used ESM1b ^30,31^. Given the limited number of labeled variants, we conducted leave-one-out calibration experiments, comparing the single-tool calibration of each DMS score to P-KNN jointly calibrating the same DMS together with ESM1b (Supplementary Methods, Integration of different evidence source).

Compared to single-tool calibration of DMS_domneg or ESM1b, P-KNN integrating both resulted in stronger evidence (mean LLR 6.01 and 4.62 for pathogenic and benign variants, compared to 5.11 and -0.01 for DMS_domneg; **Figure 6A**). P-KNN also exhibited good calibration (**Figure 6B**). In contrast, combining the evidence from DMS_domneg and ESM1b by simply summing their LLRs (according to present guidelines) resulted in pronounced overestimation of posterior probabilities (**Figure 6C**). It is likely that the miscalibration of LLR summation is driven by their statistical dependence, as the mutual information between DMS_domneg and ESM1b scores conditioned on pathogenicity was 0.057 (p=0.082) (Supplementary Methods, Conditional mutual information) ^32^. These findings suggest that, while categorized as different evidence sources (experimental vs. computational), DMS_domneg and ESM1b scores should be integrated using joint calibration.

**Figure 6.**
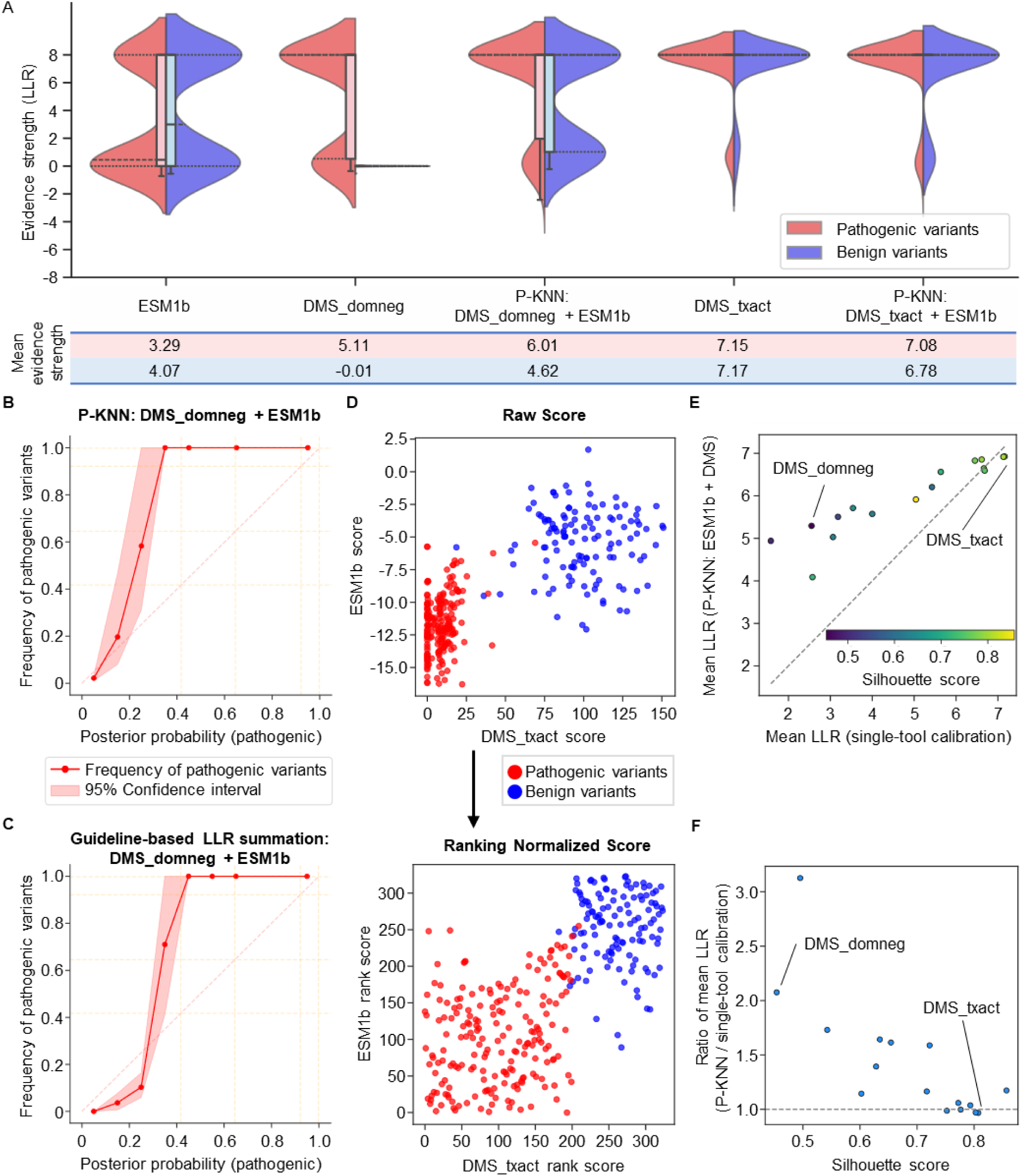
Integrating experimental and computational evidence for TP53 variant pathogenicity. **(A)** Evidence strength (log likelihood ratio) obtained by i) single-tool calibration of ESM1b, ii) single-tool calibration of a DMS assay assessing dominant negative effect (DMS_domneg), iii) P-KNN applied over ESM1b and DMS_domneg, iv) single-tool calibration of a DMS assay assessing transcriptional activity (DMS_txact), and v) P-KNN applied over ESM1b and DMS_txact. **(B)** Calibration curve for P-KNN integrating ESM1b and DMS_domneg. **(C)** Calibration curve for summing the LLRs from DMS_domneg and ESM1b (according to present guidelines). **(D)** DMS_txact and ESM1b scores across pathogenic and benign *TP53* variants, before and after ranking normalization. **(E)** Mean LLR across 16 DMS assays using single-tool calibration vs. P-KNN joint calibration with ESM1b. **(F)** The Silhouette scores of 16 DMS assays vs. mean LLR of P-KNN relative to single-tool calibration of each assay.

### Scores with near-perfect separation of pathogenic and benign variants may outperform P-KNN

For DMS_txact, single-tool calibration provides stronger evidence than P-KNN integrating DMS_txact and ESM1b (mean LLR 7.15 and 7.17 for pathogenic and benign variants, compared to 7.08 and 6.78 for P-KNN; Figure 6A). The slight reduction in evidence strength from P-KNN in this case might be explained by the fact that DMS_txact already separates pathogenic and benign variants almost perfectly (**Figure 6D**). However, the ranking normalization used in P-KNN brings the pathogenic and benign variants closer together (Figure 6D), reducing the purity of nearest-neighbor clusters near the boundary between two classes. As a result, variants located near this boundary obtain weaker evidence from P-KNN.

Altogether, P-KNN outperformed 12 of the 16 *TP53* DMS assays when integrating them with ESM1b predictions compared to individual calibrations of the same assays (**Figure 6E**). The only exceptions were 4 assays that already achieved very high evidence strength as a result of near-perfect separation of pathogenic and benign variants. To identify those cases where single-tool calibration may be preferable, we explored the Silhouette score as a simple metric of separation between the two classes (Supplementary Methods, Silhouette score) ^33^. We found that whenever the Silhouette score of an assay fell below 0.62, the score of a hypothetical predictor with perfect ranking, P-KNN consistently provided stronger evidence than the assay alone (**Figure 6F**). In contrast, assays with a Silhouette score above this threshold may perform better without P-KNN integration with ESM1b.

## DISCUSSION

This study introduces P-KNN, a joint calibration framework designed to integrate pathogenicity predictions from any set of tools in accordance with the ClinGen Bayesian framework and ACMG/AMP guidelines ^1–3^. P-KNN improves both the evidence strength and reliability of the calibrated posterior probabilities compared to single-tool calibration of individual predictors (Figure 3) and meta-predictors (Figure 4). It can also seamlessly incorporate newly developed tools, enabling dynamic and up-to-date variant classification in a highly robust manner (Figure 5). To our knowledge, this is the first framework for jointly calibrating multiple pathogenicity prediction tools.

The improvement in evidence strength achieved by P-KNN can be attributed to each prediction tool capturing distinct aspects of variant pathogenicity. When multiple tools are jointly calibrated, their combined predictions allow for better discrimination between pathogenic and benign variants. Consequently, the local neighborhoods identified in the multidimensional space of prediction scores tend to be more homogenous than neighborhoods defined by a single tool alone (Figure 1A-B). The increased homogeneity in variant pathogenicity directly translates to stronger Bayesian evidence.

P-KNN shows the greatest improvement in evidence strength for benign variants (Figure 3A, 4A, 5A). This pattern likely arises because the best-performing single tools, including AlphaMissense ^24^, REVEL ^20^, BayesDel ^19^, MutPred2 ^21^, and VEST4 ^22^, all tend to assign much stronger evidence for pathogenic variants. However, because each tool correctly classifies a different set of variants, P-KNN achieves more balanced evidence across pathogenic and benign variants.

In addition to enhancing evidence strength, P-KNN also produces better-calibrated posterior probabilities (Figure 3B–J). Our analysis demonstrates that miscalibration in single tools often arises from distribution shifts in the scores that the tool assigns to pathogenic and benign variants between the calibration and test datasets. In contrast, P-KNN mitigates this issue by leveraging a multidimensional score space across multiple tools. Because each tool exhibits its own unique distribution shift, the bias tends to average out in the joint score space. This integration acts like a signal-to-noise enhancement, where inconsistent biases cancel out while consistent patterns are reinforced, leading to overall stronger signal and more accurate calibration.

In addition to jointly calibrating multiple tools, there are three additional minor differences between P-KNN and the original single-tool calibration framework. First, while the standard single-tool calibration framework forces the mapping of raw scores to posterior probabilities to be non-decreasing, P-KNN omits this step. As a result, P-KNN is better suited to capture local variations in pathogenicity probabilities across the high-dimensional score space. However, this is not the primary reason for the stronger evidence from P-KNN. Even when the monotonicity step is removed from the standard single-tool calibration, their performance does not change much. Second, P-KNN normalizes each tool’s scores to ensure comparability across tools. As a result, when the original scaling of the score distribution provides an exceptional degree of separation between pathogenic and benign variants, as in the case of some experimental measurements (Figure 6D), ranking normalization may reduce the homogeneity of local neighborhoods and thereby diminish evidence strength. Third, the original implementation of the single-tool calibration framework gradually expands a score window around the query variant until it includes a sufficient number of variants from the calibration dataset. In contrast, P-KNN directly computes all distances and selects an exact number of nearest neighbors. In practice, this modification had minimal impact on the results while substantially reducing computational cost.

Our findings suggest that even distinct evidence sources may exhibit statistical dependency, which can be measured by mutual information conditioned on pathogenicity. When the conditional mutual information between sources of evidence is non-negligible, directly summing their calibrated evidence under the ACMG/AMP Bayesian framework may lead to inflated pathogenicity probabilities (Figure 6C) ^2,3,32^. In the specific example we studied, miscalibration is specifically observed for the lower posterior probabilities (below 0.3), suggesting that simply capping the summed evidence will not solve the problem. In such cases, joint calibration methods like P-KNN may be more reliable. Additional research is needed to determine when and how to apply each integration strategy, particularly when working with limited labeled data (as with many experimental assays).

In rare cases where functional assays already achieve close to perfect separation, integration of additional predictors with P-KNN may slightly degrade performance (Figure 6). The Silhouette score, which measures the overall separation between pathogenic and benign variants, may help identify such cases ^33^. However, more comprehensive studies of diverse datasets and genes beyond *TP53* are required to generalize the robustness of this metric. Until such studies are available, we recommend using P-KNN in all cases. While the inclusion of additional tools can in rare cases degrade performance (Figure 6A), these reductions in evidence strength are very minor, and all our analyses suggest a very high level of robustness for P-KNN, even when lower-performing tools are included.

The success of calibration critically depends on careful construction of the calibration dataset, and the most important factor is the avoidance of data contamination. Data contamination most commonly arises when some of the underlying tools have been trained on clinically annotated variants that also appear in the calibration datasets, leading to inflated posterior probabilities (Figure 2H-I). To prevent this, variants contained in the training data of any of the underlying tools must be excluded. In particular, if a prediction tool has exhausted nearly all labeled variants during training, it becomes challenging to include that tool in P-KNN (or any other calibration framework). A more subtle form of data contamination may arise in the future with the implementation of the ACMG/AMP Bayesian framework, as variants receiving strong evidence from computational tools may later end up in calibration datasets ^8^. To mitigate this, future reference databases should include information on the exact evidence sources that informed the classifications for each variant, thereby enabling the detection and management of such feedback loops.

Because P-KNN calculates distances between each pair of query and calibration variant, it requires on-the-fly computation for each query variant, unlike single-tool calibration which establishes fixed score thresholds that enable direct lookup. To enable real-time use of P-KNN, we offer a GPU-accelerated implementation capable of calibrating approximately 250 query variants per minute (over 27 tools, ∼11K calibration variants and ∼360K regularization variants). We also provide a complete catalog of precomputed posterior probabilities and LLR scores from P-KNN applied over a set of 27 state-of-the-art prediction tools, for all possible missense variants in the human genome (see Implementation and code availability, Availability of data and materials). However, given the high flexibility and robustness of P-KNN to any set of tools, and given the pace of improvement in variant pathogenicity prediction, we recommend reapplying it as newer tools become available.

## CONCLUSIONS

P-KNN enables joint calibration of any combination of pathogenicity prediction tools within a single, unified framework. By leveraging the unique strengths of each underlying tool, it produces more robust and stronger evidence than calibrated individual predictors or supervised meta-predictors, while seamlessly incorporating new predictors as they emerge.

## LIST OF ABBREVIATIONS

P-KNN: Pathogenicity K-Nearest Neighbors
ACMG: American College of Medical Genetics and Genomics
AMP: Association for Molecular Pathology
VUS: variant of uncertain significance
LLR: log-likelihood ratio
DMS: Deep mutational scanning assay
DMS_domneg: a DMS assessing dominant-negative effects
DMS_txact: a DMS measuring transcriptional activity

## DECLARATIONS

### Ethics approval and consent to participate

Not applicable

### Consent for publication

Not applicable

### Availability of data and materials

All datasets used in this study are publicly available. The P-KNN command-line tool and the full source code used to generate the study results are available at https://github.com/Brandes-Lab/P-KNN. The ClinGen datasets can be accessed at https://zenodo.org/records/8347415. The precomputed P-KNN scores for all possible missense variants in the human genome, as well as the default calibration and regularization dataset for the command line tool are hosted at https://huggingface.co/datasets/brandeslab/P-KNN. The ClinVar dataset was downloaded from https://ftp.ncbi.nlm.nih.gov/pub/clinvar/tab_delimited/, and deep mutational scanning assay data for TP53 were obtained from https://mavedb.org/.

## Authors’ contributions

PL and NB jointly conceived and designed the study. PL was responsible for data acquisition and analysis. Interpretation of the results was performed collaboratively by PL and NB. PL drafted the manuscript, and NB provided substantive revisions.

## Acknowledgements

Not applicable

