## Supplementary Material for "P-KNN: Maximizing variant classification evidence through joint calibration of multiple pathogenicity prediction tools"

### SUPPLEMENTARY FIGURES

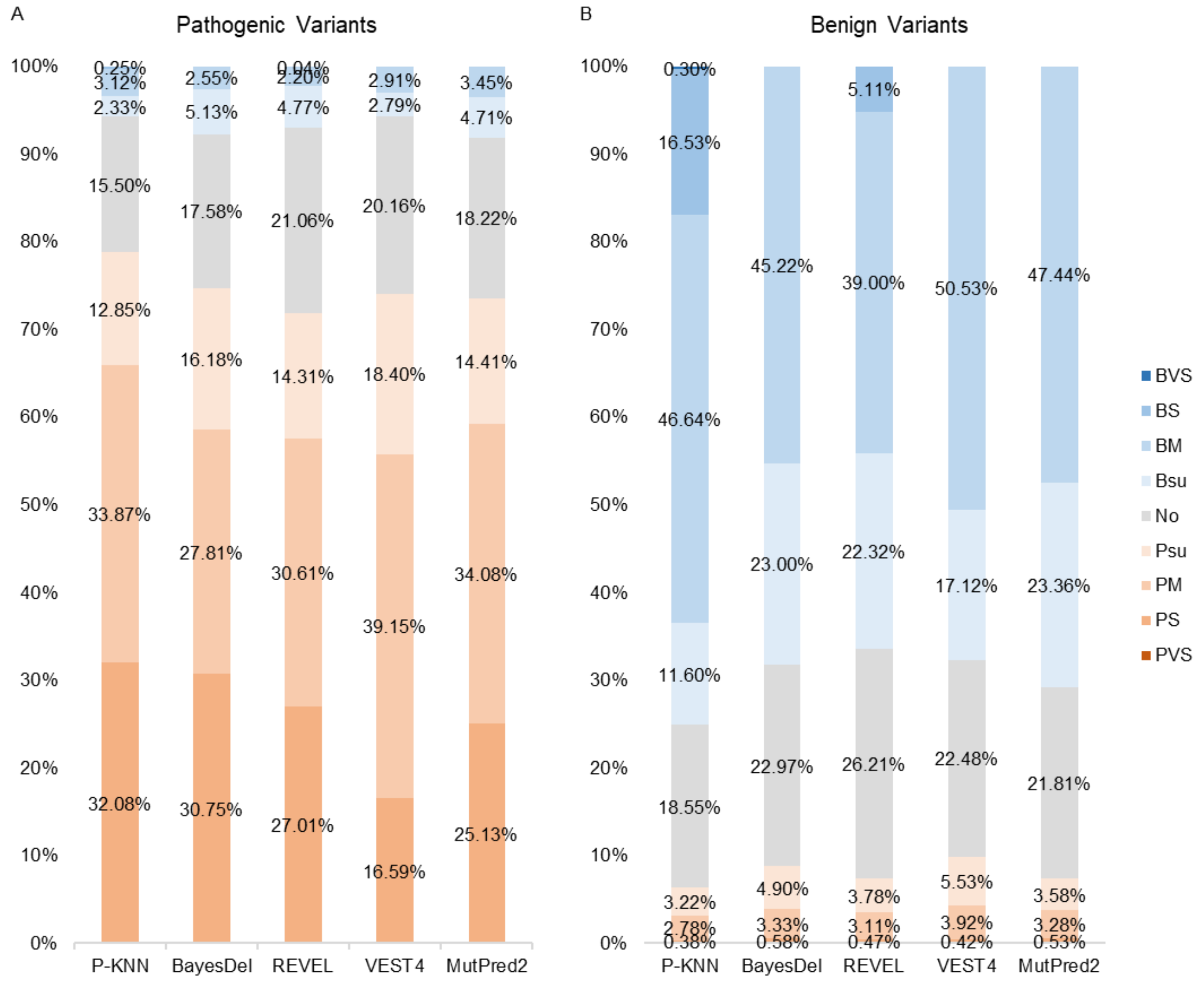


**Supplementary Figure 1. Evidence levels from P-KNN and 4 single tools.**

Rates of the different evidence levels defined by the ACMG/AMP guidelines obtained by P-KNN (integrating 13 tools) and 4 single tools calibrated individually across **(A)** pathogenic and **(B)** benign variants from ClinVar. PVS: very strong evidence (pathogenic); PS: strong evidence (pathogenic); PM: moderate evidence (pathogenic); PSu: supporting evidence (pathogenic); BVS: very strong evidence (benign); BS: strong evidence (benign); BM: moderate evidence (benign); BSu: supporting evidence (benign).

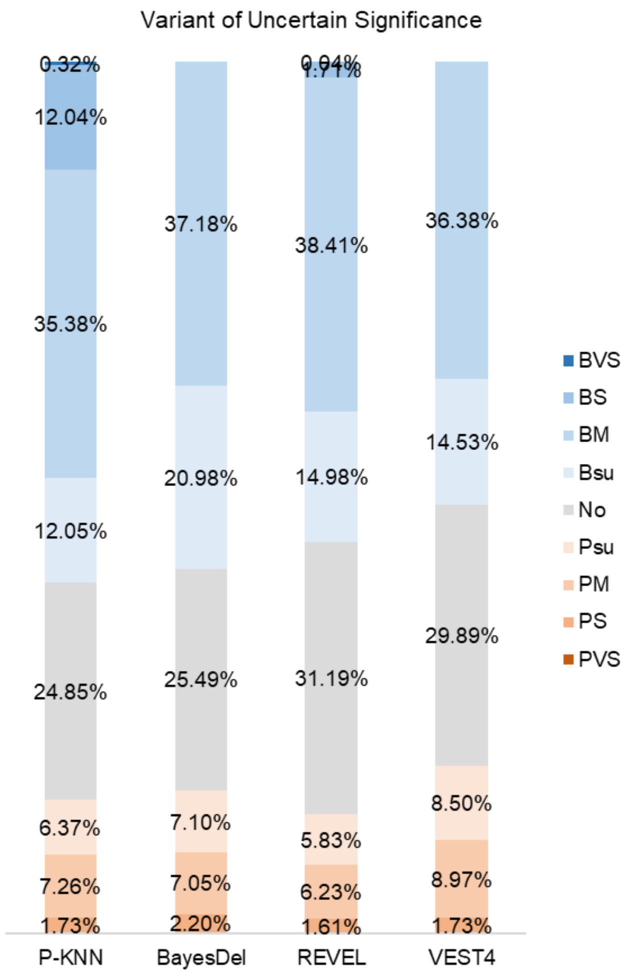


**Supplementary Figure 2. Evidence levels for variants of uncertain significance.**

Rates of the different evidence levels assigned to VUS, similar to Supplementary Figure 1. Here P-KNN includes only 11 underlying tools with available scores (see text).

### SUPPLEMENTARY TABLES

#### Supplementary table 1. Computational pathogenicity prediction tools included from different version of dbNSFP

Variant pathogenicity prediction tools included from dbNSFP v3.0, v3.5, v4.2, and v5.2 after excluding tools with overlapping training data with calibration dataset. (Tools newly added are shown in **bold**. Tools removed from dbNSFP are indicated with ~~double strikethrough~~.)

| | dbNSFP v3.0 (August 2015) | dbNSFP v3.5 (August 2017) | dbNSFP v4.2 (April 2021) | dbNSFP v5.2 (July 2025). |
|---|---|---|---|---|
| Tools inclu-ded | SIFT[1], PolyPhen-2[2] (HDIV, HVAR), LRT[3], MutationTaster[4], MutationAssessor[5], FATHMM[6], PROVEAN[7], VEST3[8], CADDv1.3[9], DANN[10], integrated fitCons[11], GERP++[12], SiPhy[13], two phyloP[14] (7 way mammalian, 20 way vertebrate), two phastCons[15] (7 way mammalian, 20 way vertebrate) | SIFT[1], PolyPhen-2[2] (HDIV, HVAR), LRT[3], MutationTaster[4], MutationAssessor[5], FATHMM[6], PROVEAN[7], VEST3[8], CADDv1.3[9], DANN[10], integrated fitCons[11], GERP++[12], SiPhy[13], two **phyloP**[14] **(20 way mammalian, 100 way vertebrate)**, two **phastCons**[15] **(20 way mammalian, 100 way vertebrate)**, **REVEL**[16], **MutPred**[17], **Eigen (Principal component)**[18], **GenoCanyon**[19] | SIFT[1], **SIFT4G**[20], PolyPhen-2[2] (HDIV, HVAR), LRT[3], MutationTaster[4], MutationAssessor[5], FATHMM[6], PROVEAN[7], **VEST4**[21], **CADDv1.6**[22], DANN[10], integrated fitCons[11], GERP++[12], SiPhy[13], two **phyloP**[14] (**30 way mammalian**, 100 way vertebrate), two **phastCons**[15] (**30 way mammalian**, 100 way vertebrate), REVEL[16], MutPred[17], Eigen (Principal component)[18], GenoCanyon[19], | SIFT[1], SIFT4G[20], PolyPhen-2[2] (HDIV, HVAR), ~~LRT[3]~~, ~~MutationTaster[4],~~ MutationAssessor[5], ~~FATHMM[6],~~ PROVEAN[7], VEST4[21], **CADDv1.7**[27], DANN[10], ~~integrated fitCons[11],~~ GERP++[12], **GERP 91 mammals,** ~~SiPhy[13],~~ three **phyloP**[14] (**17 way primate**, **470 way mammalian**, 100 way vertebrate), three **phastCons**[15] (**17 way primate**, **470 way mammalian**, 100 way vertebrate), REVEL[16], |

| | | | | |
|---|---|---|---|---|
| | | | **MPC**[23], **PrimateAI**[24], **BayesDel (no allele frequency)**[25], **LIST-S2**[26] | **MutPred2**[28], Eigen (Principal component)[18], ~~GenoCanyon[19],~~ MPC[23], PrimateAI[24], BayesDel (no allele frequency)[25], LIST-S2[26], **ESM1b**[29,30], **AlphaMissense**[31], **bStatistic**[32] |
| Tools exclu-ded | FATHMM-MKL[33], MetaSVM[34], MetaLR[34] | FATHMM-MKL[33], MetaSVM[34], MetaLR[34], **M-CAP**[35] | FATHMM-MKL[33], MetaSVM[34], MetaLR[34], M-CAP[35], **MetaRNN**[36], **MVP**[37], **DEOGEN2**[38], **ClinPred**[39] | ~~FATHMM-MKL[33],~~ MetaSVM[34], MetaLR[34], M-CAP[35], MetaRNN[36], MVP[37], DEOGEN2[38], ClinPred[39], FATHMM-XF[40], gMVP[41], VARITY_R[42], **MutationTaster2021**[4] **PHACTboost**[43], **MutFormer**[44], **MutScore**[45], |

### SUPPLEMENTARY METHODS

#### Mutual information scaling

To estimate the mutual information between the discrete pathogenicity label and the continuous score of a given tool, we employed a non-parametric KNN estimator specifically designed for discrete-continuous variable pairs.[46,47] Each genetic variant in the calibration dataset is represented as a pair $(label_i, score_i)$, where $label_i$ denotes the pathogenicity label and $score_i$ is the corresponding score from the target tool. For each variant, the algorithm computes a distance $d_i$, defined as the score difference between that variant and its $k$th nearest neighbor among variants with the same $label_i$. It then counts the total number of genetic variants $m_i$ within $d_i$ in the full dataset, regardless of label. Let $N$ denote the total number of samples, and $N_{label_i}$ the total number of samples with $label_i$, the contribution of each sample to the mutual information estimate is given by: $I_i = \psi(N) - \psi(N_{label_i}) + \psi(k) - \psi(m_i)$ where $\psi(\cdot)$ denotes the digamma function.[47] The overall mutual information is then obtained by averaging $I_i$ across all samples. Following the default setting in Scikit-learn, $k$ is set to 3.

#### Conditioned mutual information

The ACMG/AMP guidelines categorize variant classification evidence into distinct sources, and when multiple sources are combined, their evidence strengths are typically summed.[48,49] However, this summation assumes independence between sources. To determine whether two evidence sources can be directly combined or require joint calibration, we evaluate the conditional mutual information (CMI) between their scores, conditioned on the variant's pathogenicity label within the calibration dataset.[50] The CMI is computed as a weighted average of the mutual information within each class:

$$CMI = P_{pathogenic} * I(Score_A; Score_B|Pathogenic) + P_{benign} * I(Score_A; Score_B|Benign)$$

where $P_{pathogenic}$ and $P_{benign}$ denote the proportions of pathogenic and benign variants in the calibration dataset, respectively. Mutual information $I(\cdot;\cdot|\cdot)$ is estimated using only variants within the corresponding label group. To assess the statistical significance of the observed CMI, we performed a stratified permutation test in which scores from one evidence source were randomly permuted within each pathogenicity class and re-calculated CMI. This process was repeated 1,000 times to generate a null distribution of CMI values, and the one-sided p-value was calculated as the proportion of permuted CMIs greater than or equal to the observed CMI.

**Silhouette score**

We use the Silhouette score to evaluate whether a single pathogenicity prediction tool can perfectly separate pathogenic from benign variants. Variants in the calibration dataset are grouped into two clusters based on their labels, and the Silhouette score quantifies how well the tool's scores separate these clusters by measuring, for each variant, the relative distance to variants in the opposite cluster versus those within the same cluster.[51] Higher values indicate better separation. Based on empirical analysis of a hypothetical predictor with perfect ranking (area under ROC curve 1.0), the maximum Silhouette score achievable using rank-transformed scores is 0.62. Tools exceeding this threshold are considered capable of near-perfect separation.

**Code implementation**

For KNN imputation, mutual-information scaling, conditional mutual information, and Silhouette score, we employed Scikit-learn's KNNImputer, mutual_info_classif, mutual_info_regression and silhouette_score, respectively [52]. We also implemented a custom PyTorch-based KNNImputer that replicates the algorithm used in Scikit-learn.

### Evaluation with simulated tools.

#### *Generation of simulated pathogenicity prediction tools*

To generate simulated pathogenicity prediction tools, we first created a dataset of 14,500 genetic variants, each with 17 randomly sampled features ranging between 0 and 1. These features were processed through a randomly parameterized three-layer multilayer perceptron, yielding a single scalar output per variant. Variants with output values in the upper half of the distribution were labeled as pathogenic, while those in the lower half were labeled as benign. We then introduced 1% Gaussian noise to the 17 original features and partitioned the resulting variants into four subsets: 50,000 variants for model training, 10,000 for calibration, 35,000 for regularization, and 50,000 for testing.

To simulate pathogenicity prediction tools, we removed two features from the model training dataset and randomly selected 8 to 12 of the remaining features to train each tool. Using Scikit-learn, we trained a total of 50 tools based on three model types: Gaussian Naive Bayes, Random Forest Classifier, and Logistic Regression.[52] For the Random Forest models, the minimum number of samples required to split an internal node was set to 10; all other hyperparameters were kept at their default settings.

#### *Evaluation of score standardization and calibration dataset contamination*

Using the scores generated from the 50 simulated tools, we evaluated four score preprocessing pipelines by varying two factors: (1) rank-based normalization versus z-score normalization, and (2) mutual-information scaling applied or not applied. We assessed the effect of each pipeline under two conditions: (a) different total numbers of tools being jointly calibrated, and (b) a fixed total 15 tools with varying proportions of highly informative tools.

To evaluate handling of missing values in the calibration dataset, we compared KNN imputation against the alternative of excluding variants with missing values. Performance was assessed under varying maximum percentages of missing values per tool (each tool assigned a random missing rate between 0 and the maximum up to 40%; missing values inserted randomly) and under four different missingness patterns: random, or biased toward high, low, or mid-range scores, with a maximum missing rate of 25%.

After identifying the optimal preprocessing pipeline, we tested its performance using a fixed set of 15 tools with up to 25% missing values per tool. Additionally, we examined the impact of expanding the calibration dataset by incorporating multiples of the model training dataset, evaluating how calibration dataset contamination with model training data affects overall P-KNN performance.

### Evaluation with established tools on real clinical data

#### *Evaluation on labeled pathogenic and benign variants*

To evaluate the performance of P-KNN relative to single-tool calibration developed by ClinGen, we utilized three datasets curated by the ClinGen.[53] The ClinVar 2019 dataset, which serves as the calibration dataset, includes variants classified as pathogenic, likely pathogenic, likely benign, or benign prior to December 2019 in ClinVar database.[54] The ClinVar 2020 dataset serves as the test set and contains variants classified as pathogenic, likely pathogenic, likely benign, or benign, submitted to ClinVar in 2020. The gnomAD dataset, used as regularization dataset, includes all variants in gnomADv2 after excluding those in ClinVar 2019.[55] All three datasets were subject to the same preprocessing criteria, including allele frequency < 0.01, exclusion of variants located in genes without known pathogenic variants, and best-effort removal of any variants used in training the 13 tools. Each datasets includes 13 prediction tools, including two meta-predictors: REVEL[16] and BayesDel (no allele frequency)[25], and eleven individual-predictors: MutPred2[28], VEST4[21], CADDv1.6[9], PolyPhen-2 (HumVar model)[2], FATHMM[6], MPC[23], PrimateAI[24], SIFT[1], Evolutionary Action[56], GERP++[12], and PhyloP (100 way vertebrate)[14]. Using P-KNN, we joint calibration scores from all 13 tools and compared the performance with those obtained through single-tool calibrations of each tool. The comparison of evidence strength was performed by Wilcoxon signed-rank test.

***Evaluation on variants of uncertain significance***

To assess the utility of P-KNN in supporting variant classification for uncertain cases, we evaluated its performance on missense variants labeled as variants of uncertain significance (VUS) in ClinVar (Release 2025.03, tab delimited format) with a review status of at least one gold star.[54] Consistent with ClinGen criteria, variants with an allele frequency > 0.01 in gnomAD v2.1 were excluded.[55]

Although the full ClinGen dataset includes 13 prediction tools, only 11 of them were available in the dbNSFP we used as the source for prediction scores for VUS [57] We annotated each variants with 11 tools that were also used in the ClinGen dataset. Ten of these tools were sourced from dbNSFP v5.1: REVEL[16], BayesDel (no allele frequency)[25], VEST4[21], PolyPhen-2 (HumVar model)[2], FATHMM[6], MPC[23], PrimateAI[24], SIFT[1], GERP++[12], and PhyloP (100 way vertebrate)[14]. One tool CADDv1.6[9] was obtained from dbNSFP v4.2 via ANNOVAR.[57,58] Using P-KNN, we jointly calibrated the scores from these 11 tools and compared the resulting evidence strengths with those obtained from single-tool calibrations of the underlying tools. The comparison of evidence strength was performed by Wilcoxon signed-rank test on the absolute values of log likelihood ratios.

#### P-KNN vs. meta-predictors

We utilized the genetic variants in ClinVar 2019, ClinVar 2020, and gnomAD datasets provided in the ClinGen calibration study, focusing on the two included meta-predictors: REVEL and BayesDel.[16,25,53] All variants were annotated with REVEL's component pathogenicity prediction tools using ANNOVAR and dbNSFP.[57-59] (BayesDel was trained on a subset of these component tools.) Specifically, the annotations included SIFT[1], PolyPhen-2[2] (HDIV and HVAR), LRT[3], MutationTaster[4], MutationAssessor[5], FATHMM[6], VEST3[8], GERP++[12], and SiPhy[13], as well as three phyloP[14] (46 way primate, 46 way placental, 100 way vertebrate) and three phastCons[15] (46 way primate, 46 way placental, 100 way vertebrate) from dbNSFP v2.7. In addition, PROVEAN[7] annotations were obtained from dbNSFP v2.9 and MutPred[17] from dbNSFP v4.7. Since a single variant can yield different scores across transcripts, we selected all transcripts that produce a missense effect at that genomic coordinate and averaged their scores. We compared the single-tool calibration of REVEL and BayesDel, and three P-KNN: (1) integration of REVEL's component tools only, (2) REVEL combined with its component tools, and (3) integration of REVEL, BayesDel, and REVEL's component tools.

#### P-KNN evolution with new tools

We utilized the genetic variants in ClinVar 2019, ClinVar 2020, and gnomAD datasets from the ClinGen calibration study to evaluate how the performance of P-KNN evolves with the inclusion of newly developed prediction tools over time.[53] All variants were annotated using four distinct sets of variant pathogenicity prediction tools derived from different versions of the dbNSFP database: v3.0 (released August 2015), v3.5 (released August 2017), v4.2 (released April 2021), and v4.7 (released March 2024).[57,60] Since a single variant can yield different scores across transcripts, we selected all transcripts that produce a missense effect at that genomic coordinate and averaged their scores. To prevent data leakage, we excluded any tools with known or potential overlap in training data with the calibration dataset. The remaining tools from each dbNSFP version formed tool sets containing 18, 22, 27, and 33 tools, respectively (Supplementary Table 1). We applied P-KNN to each set to evaluate how performance changes as newer tools are incorporated over time.

**Integration of different evidence source**

From the March 2025 release of ClinVar, we selected all variants in the *TP53* gene with a minimum annotation quality of one star and classified as pathogenic, likely pathogenic, benign, *or* likely benign.[54] Variants with an allele frequency greater than 0.01 in gnomAD v2.1 were excluded.[55] The remaining variants were annotated with scores from ESM1b[29,30] and 16 Deep mutational scanning (DMS) assays obtained from MaveDB, focusing on one measuring transcriptional activity and the other assessing dominant negative effect.[61-63]

We performed single-tool calibration for each of the 17 scores (ESM1b and the 16 DMS assays), as well as P-KNN joint calibration of the each DMS assay with ESM1b. In addition, we tested the direct summation of their evidence strengths according to present guidelines. Due to the limited sample size, we employed a leave-one-out strategy, treating one variant as the query variant while using the remaining variants as the calibration dataset. The full set of variants, including the query variant, was used as the regularization dataset. The minimum number of calibration variants required within the neighborhood $N_{calibration}$ was set to 10. The prior pathogenicity probability was set to 0.22, following a prior research for calibrating DMS-based evidence.[64] To assess whether these scores should be jointly calibrated, we evaluated the conditional mutual information between scores given the variant label (Supplementary method, Conditioned mutual information), as well as their Silhouette scores (Supplementary method, Silhouette score), to determine the degree of independence and separability among the tools.
